# Newly isolated sporopollenin microcages from *Cedrus libani* and *Pinus nigra* for controlled delivery of Oxaliplatin

**DOI:** 10.1101/2020.10.19.345157

**Authors:** Muhammad Mujtaba, Bahar Akyauz Yılmaz, Demet Cansaran-Duman, Lalehan Akyuz, Sevcan Yangın, Murat Kaya, Talip Çeter, Khalid Mahmood Khawar

**Affiliations:** Institute of Biotechnology, Ankara University, Ankara, 06110, Turkey; Department of Biotechnology and Molecular Biology, Faculty of Science and Letters, Aksaray University, 68100, Aksaray, Turkey; Aksaray University, Technical Vocational School, Department of Chemistry Technology, 68100, Aksaray, Turkey; Kastamonu University, Faculty of Arts and Sciences, Department of Biology, 37100 Kastamonu, Turkey; Department of Field Crops, Faculty of Agriculture, Ankara University, 06100 Ankara, Turkey

**Keywords:** xCELLigence, real-time cell analyzer, controlled-drug release, sporopollenin

## Abstract

Sporopollenin-mediated controlled drug delivery has been studied extensively owing to its physicochemical and biological charachteristics. In the present study, sporopollenin was successfully extracted from pollen grains of *C. libani* and *P. nigra* followed by the loading of a commonly known anticancer drug Oxaliplatin. Both the drug loading and physicochemical features were confirmed by using light microscopy, FT-IR, SEM and TGA. For the first time, real-time cell analyzer system, xCELLigence, was employed to record the Oxaliplatin-loaded and sporopollenin-mediated cell death (CaCo-2 and Vero cells) in real time. Both the assays confirmed the slow release of Oxaliplatin from sporopollenin for around 40–45 h. The expression of MYC and *FOXO-3* genes significantly increased in CaCo2 cell and decreased non-cancerous Vero cell confirming that sporopollenin-mediated controlled release of Oxaliplatin was promoting apoptosis cell death preventing the spread of its negative effects to nearby healthy cells. All the results suggested that *C. libani* and *P. nigra* could be suitable candidates for slow delivery of drugs.

## Introduction

Designing and producing drug carrier systems for slow and targeted release to prevent premature burst release of the content in the bloodstream represent an ever-evolving research and application area in biomedical science [1–4]. In recent years, with the advancements in nanotechnology, huge efforts have been made to design carrier systems exhibiting the desired structural and chemical properties [5]. Nano/micro vehicles tested so far for the controlled release of anticancer drugs include liposomes, water-soluble polymers, dendrimers, vesicles [6], polymeric nanoparticles [7], and some inorganic materials [8–10]. All the tested drug carriers have their pros and cons. For example, the encapsulation capacity of liposomes and nanoparticles has been reported to be around 10% [11–13]. Besides, obtaining nanocarriers with the desired geometry and functional properties requires expensive processes and a longer time [14]. In this regard, the quest has been underway for producing alternative macro/nano/microcarriers that offer several similar properties such as being non-toxic, non-reactive with load, economical, and biodegradable. In nature, a variety of biomaterials are already present that have the application capabilities in different fields. These natural materials have evolved over a longer period, thus ensuring their own fidelity and physicochemical properties. Among many examples, pollen comes with excellent features such as excellent elasticity, size uniformity, homogeneity in pore sizes, physically and chemically resistant, UV shielding ability, and antioxidant ability [15, 16].

Since the last decade, researchers have actively focused on the development and modification of plant pollen as a delivery vehicle for many active ingredients such as drugs [17–19] and active compounds [20]. Considering the latest research related to the utilization of sporopollenin for the controlled delivery of anticancer drugs, researchers are conducting *in-vitro* release assays using different buffer solutions to simulate the intestine and stomach environment and to investigate the release effect of the drug from carrier [21–23]. However, these assays have not been conducted using novel real-time cell analyzer tools, such as xCELLigence, to get a more accurate picture of the controlled release in cancer cells. In order to bridge this gap in the literature, besides the conventional release assays (simulated pH systems), the current study is mainly designed to get an inclusive insight of the sporopollenin-based release in cancer cells by using a recently developed tool known as real-time cell analyzer. Most of the anticancer drugs exhibit problems such as poor water solubility, rapid blood clearance, low tumor selectivity, and severe side effects for healthy tissues [13]. In order to overcome these problems, several delivery systems have been tested for controlled release of anticancer drugs. As is known, in apoptosis a cell disintegrates in a rather controlled way from inside minimizing damage and disruption to the surrounding healthy cells. The resulting debris is then removed through phagocytes [24]. In necrosis, however, the abrupt disintegration of cells releases debris into the surrounding healthy cells [25]. Considering these facts, it can be assumed that the slow release of the therapeutic drug may have a direct link with cell apoptosis. In the current study, two reference genes, *c-Myc* and *FOXO3*, have been selected to assess the effect of sporopollenin-mediated slow release of Oxaliplatin on cell death by apoptosis. *c-Myc* gene plays a regulatory role in cell proliferation and cellular metabolism by inducing apoptosis [26]. The other gene, *FOXO3*, promotes apoptosis through the expression of genes responsible for cell death [27]. Thus, by determining the levels of expression of both genes, the authors have determined the possible relationship of sporopollenin to apoptosis.

In the study, the sporopollenin of *C. libani* and *P. nigra* was tested as a controlled release vehicle for a common anticancer drug known as Oxaliplatin. Pinus and Cedrus belong to the Pinaceae family. Around, 80 species of the Pinus genus are distributed from the Northern hemisphere to the Tropics [28]. Five naturally grown species are found in the authors’ country. These are 10–20 m tall, evergreen trees with two-to-three or even five needle leaves growing out of the shoots [29]. Male cones are located at the bottom of small, yellow, and young shoots; and female cones are in the green areas close to the tip. *Pinus nigra* pollen is monad, heteropolar and with bilateral symmetry. The pollen shape is equatorially oblate and its type is either vesiculate or bisaccate. *Cedrus libani* pollens are monad and heteropolar with bilateral symmetry. The pollen shape is oblate at both ends and the type is either vesiculate or bisaccate [30].

To the best of the authors’ knowledge, no study has yet been conducted on *C. libani* and *P. nigra* sporopollenin production and its utilization as a controlled drug vehicle for Oxaliplatin and the effect of the consequent slow release on cell death. Besides the conventional release measurement assay (simulated pH systems), a real-time cell analyzer model (xCELLigence) was employed for the first time to determine the slow release from sporopollenin in real time. Here, sporopollenin production and drug loading were confirmed by using analytical tools such as light microscopy, SEM, FTIR, and TGA. The release assays were conducted both in simulated pH solutions and cell culture (Caco-2 and Vero) by using a real-time cell analyzer (xCELLigence). The effects of sporopollenin-mediated slow release of Oxaliplatin on apoptosis were investigated. The cell apoptosis was analyzed by using two reference genes, known as *FOXO-3* and *c-MYC*, which had the main role in the apoptosis.

## Materials and methods

### Pollen collection

In the current study, the pollens of *Cedrus libani* and *Pinus nigra* were collected in the year 2015 from the gardens of Education Faculty and Vocational School of Kastamonu University, respectively. Male cones collected with vintage scissors were placed in clean paper bags and brought to the laboratory. They were then spread over clean sheets of paper and covered with paper to prevent contamination and allowed to dry at room temperature for three days. The dry male cones were shaken to shed pollens. The pollens thus collected were sieved (having a pore diameter of 10 μm) to be free from dust and other unwanted particles. In order to obtain pure pollens, the sieved pollens were again passed through another sieve having a pore diameter of 100 μm, thus removing larger particles. The purified pollen samples were stored in sealed falcon tubes at −20°C for further experiments.

### Chemicals

CH_3_OH, CHC1_3_, HCl, KCl, KH_2_PO_4_, NaCl, NaOH and Na_2_HPO_4_ have been purchased from Sigma-Aldrich (St. Louis, Missouri, USA). PBS (phosphate buffer saline) buffer was prepared at pH 7.4 by using Na_2_HPO_4_, KH_2_PO_4_, NaCl and KCl. Distilled water was used in all experimental steps. DMEM (Sigma, UK), Penicillin/streptomycin (Sigma, UK) and non-essential amino acid (Gibco, Grand Island), Fetal Bovine Serum (Biological Industries, USA) and PBS (Biological Industries, USA) were also used.

### Isolation of Sporopolenin

In order to obtain sporopollenin, the stored pollens were treated with acid, base and chloroform: methanol: water solution for demineralization, deproteinization and depigmentation. For demineralization, 10 g of *C. libani* and *P. nigra* pollens were treated with 4M HCl solution at 50°C for 2 h. Then the samples were filtered with Whatman filter paper having a pore size of 110 μm followed by an extensive wash with distilled water until neutral pH. For deproteinization, the samples were incubated with 4 M NaOH solution at 150°C for 6 h. Then the treated samples were filtered with Whatman filter paper and extensively washed with distilled water until neutral pH. The demineralization and deproteinization treatments were repeated four times to ensure complete removal of genetic and cellulosic materials inside the pollens. The acid- and base-treated pollen samples were then kept submerged in a chloroform: methanol: water solution (4: 2: 1 / v: v: v) at room temperature for 1 h. Finally, the samples of sporopollenin were thoroughly washed with distilled water and dried in an oven at 60°C for 48 h [17]. The samples were coded Oxaliplatin drug (*OX*), *Pinus nigra* raw pollen (PN), *Pinus nigra* sporopollenin (*PN-SP*), Oxaliplatin loaded *Pinus nigra* sporopollenin (*PN-SP-OX*), *Cedrus libani* raw pollen (*CD*), *Cedrus libani* sporopollenin (*CD-SP*), and Oxaliplatin loaded *Cedrus libani* sporopollenin (*CD-SP-OX*).

### Physicochemical analysis

#### FTIR

The spectra of Oxaliplatin drug (*OX*), *Pinus nigra* raw pollen (PN), *Pinus nigra* sporopollenin (*PN-SP*), Oxaliplatin loaded *Pinus nigra* sporopollenin (*PN-SP-OX*), *Cedrus libani* raw pollen (*CD*), *Cedrus libani* sporopollenin (*CD-SP*), Oxaliplatin loaded *Cedrus libani* Sporopollenin (*CD-SP-OX*) were analyzed in the range of 4000-650 cm-1 using an FT-IR spectrometer (Perkin Elmer 100, USA) under ambient conditions.

#### TGA

All samples were heated from 30 °C to 650 °C with a temperature change of 10 °C/min and their thermal stability, water and ash content were determined. All the thermograms were recorded with an EXSTAR S11 7300 (USA) instrument under the nitrogen atmosphere.

#### SEM

The drug, pollen and sporopollenin samples were placed on aluminum stab with double side adhesive type and it was covered with gold by Cressington, Sputter Coater 108 Auto Au-Pd Coating Machine at Kastamonu University Central Research Laboratory, under 40 mA for 30 seconds. SEM photos were taken with FEI-Quanta FEG 250 model scanning electron microscope.

### Light microscopy

The sporopollenin production and drug loading were also confirmed by analyzing the samples under a LEICA DFC425 C light microscope. The samples were analyzed in ambient conditions using glass slid.

### Loading of Oxaliplatin to sporopollenin via passive loading technique

Oxaliplatin was loaded onto *C. libani* and *P. nigra* sporopollenin through the passive loading technique used in our previous study (Mujtaba, Sargin et al. 2017). In the passive loading technique, 100 mg of Oxaliplatin was dissolved in two samples of 4 ml of distilled water each with the aid of sonication for 10min. Then 200 mg of *C. libani* and *P. nigra* sporopollenin were added to the solutions separately and the suspensions were vortexed for 15 min. Each sporopollenin-Oxaliplatin solution was then incubated at 350 rpm using a thermo-shaker for 4 h and at 4°C. Each sample was then filtered with filter paper having a pore size of 110 μm and washed twice with 3 ml of distilled water. The drug-loaded sporopollenin samples were incubated at −80°C for 30 minutes. Finally, the Oxaliplatin-loaded sporopollenin samples were dried at room temperature for 24 h and then stored at −18°C until further experiments.

### Encapsulation efficiency

Quantitative determination of Oxaliplatin loaded on *C. libani* and *P. nigra* sporopollenin was carried out by following previously reported method with minor modifications (Mundargi, Tan et al. 2016). Oxaliplatin was dissolved in different concentrations, that is, 5 μg/ ml, 10 μg / ml, 15 μg / ml, 20 μg / ml and 25 μg / ml to obtain the calibration curve. The absorbance of these solutions was measured at 255 nm using a UV-vis spectrophotometer. To evaluate the drug loading efficiency of both sporopollenin samples, 10 mg of Oxaliplatin loaded *C. libani* and *P. nigra* samples were vortexed for 10 min by adding 3 ml of PBS. Then the samples were filtered through Whatman filter paper with a pore size of 110 μm. The clear solution was collected and measured at 255 nm using UV-spectrophotometer. The following equations are used to calculate the loading efficiency:

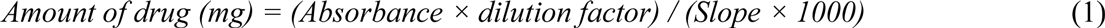

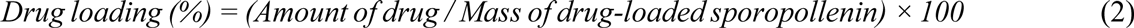

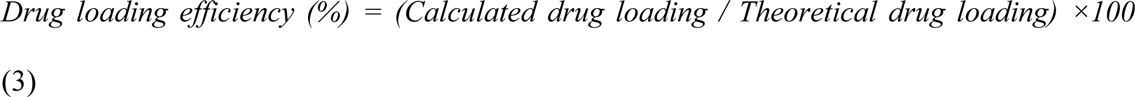

### *In vitro* release studies with Oxaliplatin-loaded microcages

*In vitro*, drug release studies of Oxaliplatin-loaded sporopollenin obtained by using the passive loading technique were carried out with PBS (pH 7.4) buffer as simulated intestinal fluid. Briefly, 10 mg of drug-loaded sporopollenin was added separately to two samples each of 5 ml PBS buffer solution. Each solution was then transferred to a dialysis bag and placed in 50 ml of PBS solution. The samples were then shaken in a water bath at 37°C and at 100 rpm for 120 h. Then, to calculate the amount of Oxaliplatin released from the sporopollenin contained in the dialysis bags into the PBS solution, 2 ml of PBS was drawn at 5^th^ min, 15^th^ min, 30^th^ min and at 1^st^, 2^nd^, 6^th^, 24^th^, 48^th^, 72^nd^ and 120^th^ h time intervals. The same amount of fresh PBS buffer solution was added back to the medium for maintaining the volume. The amount of Oxaliplatin released into the medium was measured using a *UV-VIS* spectrophotometer at 255 nm. Besides, in order to control the release of free drug, 10 mg Oxaliplatin was added to the PBS solution and *in vitro* release studies were performed by following the above procedure.

The following equations were used to calculate the cumulative percent release of Oxaliplatin:

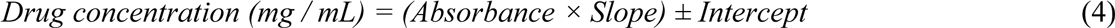

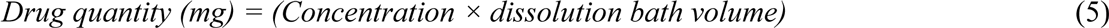

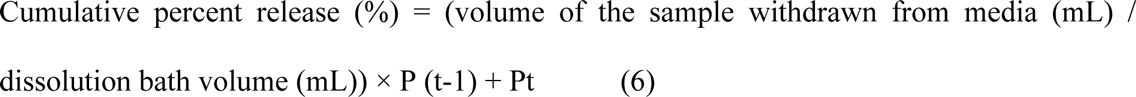

*Pt* denotes the percentage release at time *t*, *P* (*t* − 1) is the percentage release previous to *t*.

### Real-time release assay on cell lines

#### Cell culture

CaCo-2 and Vero cells were kindly provided by the Ministry of Health, Turkey Pharmaceuticals and Medical Devices Agency. The CaCo-2 cancer cell was cultured in Dulbecco’s modified Eagle’s medium (DMEM, Gibco, Grand Island, NY, USA) supplemented with 1% penicillin/streptomycin (Gibco, Grand Island, NY, USA), 1% amino acid (Gibco, Grand Island, NY, USA), 20% fetal bovine serum (Lonza, Verviers, Belgium), at 37°C in a humidified atmosphere containing 5% CO_2_. The non-cancerous cell line, Vero, was cultured in DMEM (Gibco, Grand Island, NY, USA) supplemented with 10% fetal bovine serum (Lonza, Verviers, Belgium), 1% penicillin/streptomycin (Gibco, Grand Island, NY, USA) in a 5% CO_2_ humidified incubator at 37°C.

### Anti-proliferative effect of sporopollenin-based control release (xCELLigence, RTCA)

The cell proliferation was continuously monitored by using the xCELLigence RTCA Instrument (xCELLigence RTCA, Roche, Germany) according to the manufacturer’s instructions. CaCo-2 and Vero cells seeded at a density of 5 × 10^3^ cells and 7 x 10^3^ per well in 200 μl media containing 10% FBS in a 16-well E-plate (Roche Applied Science, Germany), respectively. After 24 h, drug-loaded sporopollenin of *P. nigra*, *C. libani*, and pure Oxaliplatin (control) were added at different concentrations (in the range of 5–20 mg/ml). The E-plates were continuously monitored on an RTCA system for 120 h at 37°C with 5% CO_2_. The proliferation of examined cells was monitored every 30 min and a time-dependent cell index (CI) graph was produced by the device using the RTCA software program of the manufacturer. The baseline cell index for molecules-treated cells, compared to cancer and control cells, was calculated for at least two measurements from three independent experiments.

### Total RNA isolation and cDNA synthesis

The cells were plated at a density of 5x 10^5^ cells in a 6-well plate with a 2 ml culture medium. After keeping the drug-loaded *C. libani* and *P. nigra* sporopollenin for 24 h. in the cell culture of CaCo-2 and Vero, the cells were washed twice with PBS and total RNA was isolated with the help of Trizol Reagent. According to the manufacturer’s instructions, 1 μg RNA was reverse transcribed by oligo (dT) primers with Transcriptor High Fidelity cDNA Synthesis Kit (EUX, Germany).

### Quantitative real-time RT-PCR

Quantitative real-time RT-PCR analysis was performed using a Lightycycler 480 (Roche, Germany). Complementary DNA (10 ng), 2.1 μL of PCR-grade water, 5 μL of SYBR Green real-time PCR master mix and 1.5 μL of primer pairs were mixed in a PCR tube. PCR was performed with an initial denaturation at 95°C for 10 seconds, followed by amplification for 40 cycles, each cycle consisting of denaturation at 95°C for 5 seconds, annealing at 65 °C for 60 seconds. The primers were as follows in Table 1. The housekeeping gene, GAPDH, was used for normalization. The qRT-PCR experiments were repeated three times. The 2^−ΔΔCT^ method was used to calculate the fold change of mRNA expression level. To validate the size of amplified fragments, PCR products were separated by electrophoresis through 2.0% agarose gels and visualized with ethidium bromide.

**Table 1:**
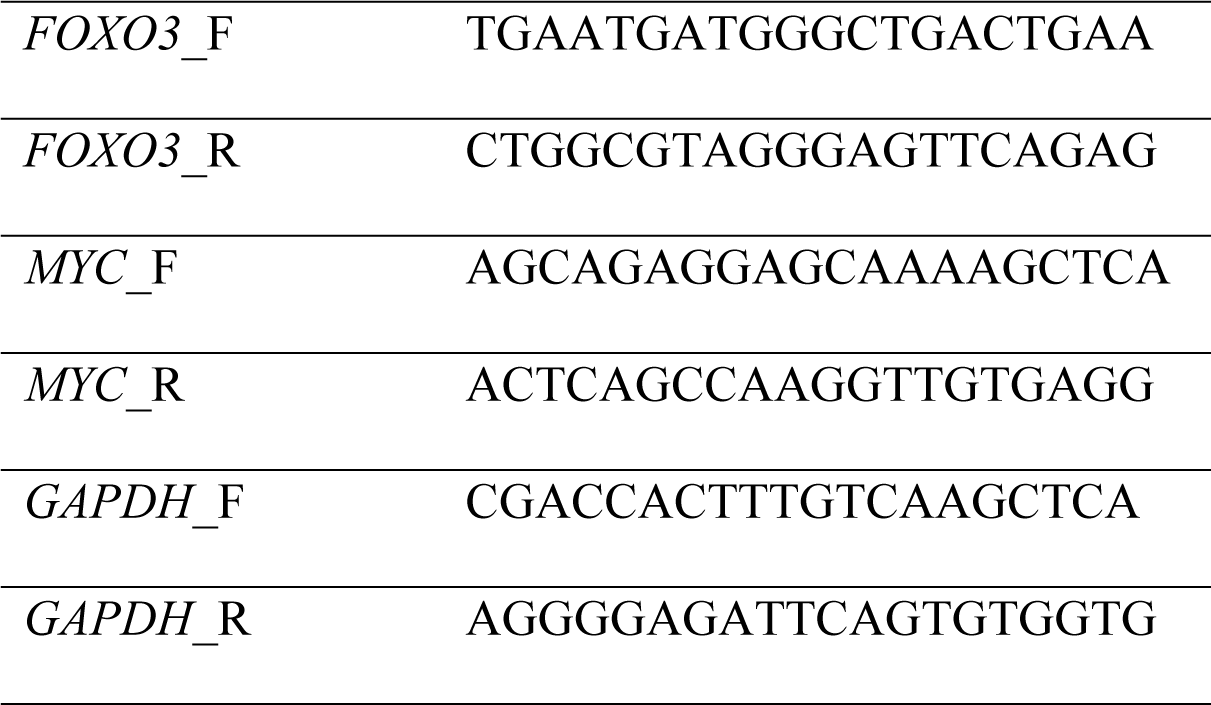
The sequences of forward (_*F*) and reverse (_*R*) primers 5'→3') of *FOXO3* and *MYC* genes.

## Results and discussions

### FT-IR

FT-IR analysis was carried out to investigate the chemical interactions between sporopollenin and drug. The FT–IR spectra of *OX, CD, CD-SP, CD-SP-OX, PN, PN-SP* and *PN-SP-OX* are given in Fig 2. In the spectrum of OX, the peaks observed at 3211.3 cm^−1^ and 3160.1 cm^−1^ were ascribed to N–H bonding. The sharp peak at 1696.3 cm^−1^ corresponded to the stretching vibrations of the carbonyl groups (C=O) of *OX*. The N-H bending peak was observed at 1609.7 and 807.82 cm^−1^. The authors also observed that the FT–IR spectra of both raw pollens, that is, *CD* and *PN* were similar. The broad peaks were observed at about 3345 cm^−1^ due to the intermolecular O-H bonding in the FT–IR spectra of raw pollens (*CD* and *PN*). Amide I and amide II linkages of the protein structure of raw pollens appeared at 1605 and 1515 cm^−1^ for both pollens. The stretching peaks of the sugar rings of polysaccharides in raw pollens were recorded as the broad peaks between 1100–1028 cm^−1^ for *CD* and *PN*.

Although a certain chemical structure of sporopollenin has not been explained until today, some research groups have identified unsaturated fatty acids, olefinic and aromatic functional groups found in sporopollenin [31]. The authors have determined that sporopollenin, which consists of C, H and O, is an organic polymer and is nitrogen-free [32, 33]. The chemical structure of sporopollenin can be different by species [33]. In the present study, the FT–IR spectra of CD-SP and PN-SP show that the chemical structures of the extracted sporopollenin are different from those of *CD* and *PN*. When the *CD* spectrum was compared with the FT–IR spectrum of the CD-SP, the new peaks were observed at 1575.6 and 1538.9 cm^−1^ due to the sporopollenin polysaccharide structures. In the PN-SP spectrum, the O-H bonding peak was shifted to 3325.2 cm^−1^ and the intensity of the peak decreased. However, a sharp peak appeared at 1713.4 cm^−1^ due to the olefinic acid structures. In the *CD-SP-OX* spectrum, it was seen that the C-H stretching of *CD-SP* shifted to a greater wave number due to the effect of the C-H stretching of drug molecules. N-H bending vibrations of Oxaliplatin were observed at 1603.6 cm^−1^. Alkane C-H bending of *OX* and C-H bending of *CD-SP* overlapped and intensified at 1378.3 cm^−1^. In the *PN-SP-OX* spectrum, the O-H band was becoming broader due to the effect of the N-H bending of *OX*. The peaks intensified in the region of 12451500 cm^−1^ because of the drug molecules. These results showed that the drug had been successfully loaded onto *CD-SP* and *PN-SP*.

### Scanning electron microscopy (SEM) and light microscopy analysis

SEM analysis was carried out to evaluate the pollen and sporopollenin structures and to better understand the drug loading mechanism. *C. libani* and *P. nigra* have monad pollens consisting of two pollen saccus which are connected to the main body called Corpus at the distal pole called Corpus. The main body of the pollen is surrounded by two wall layers, intine and exine. The pollen saccusis alveolar vesicles consist of exine (Fig 3c, g, and 1b, f). As is known, sporopollenin isolation goes through two steps, acid hydrolysis and base hydrolysis, for the removal of internal proteins and pigment structures. The same procedure was followed for the production of *CD-SP* and *PN-SP*. SEM micrographs confirmed the successful isolation of sporopollenin. Compared to the untreated pollens, the porous surface becomes visible after the removal of organic compounds through acid and base treatments. SEM micrographs revealed that acid treatment led to the structural disintegration of sporopollenin. Following the acid hydrolysis treatment, *CD* preserved its structural integrity to a large extent (85–90%), and transformed into sporopollenin capsules (cages) only with a small cleavage on the leptoma (Fig 3d and 1g). In the case of *PN*, the acid hydrolysis resulted in complete structural disintegration (90–95%) and formed structures or plaques resembling sporopollenin sheets. Only a small percentage (5–10%) of the pollens retained their original shape (cages), leaving only a cleavage on the leptoma (Fig 3h and 1c). Although the acid hydrolysis resulted in the structural deformation of *PN-SP*, the sporopollenin was porous which could help trap or adsorb the Oxaliplatin particles on its surface (Fig 3d, h). In the current study, a commonly known anticancer drug Oxaliplatin was applied in the form of a water solution. Considering the cumulative control release and drug encapsulation results, it could be assumed that the drug had been encapsulated into *CD-SP* by adsorption or direct penetration through porous walls (Fig 3e, f and 1h). *PN*, which were highly decomposed into sporopollenin plaques, formed big granules together with Oxaliplatin solution (Fig 3i, j and 1d).

**Figure 1.**
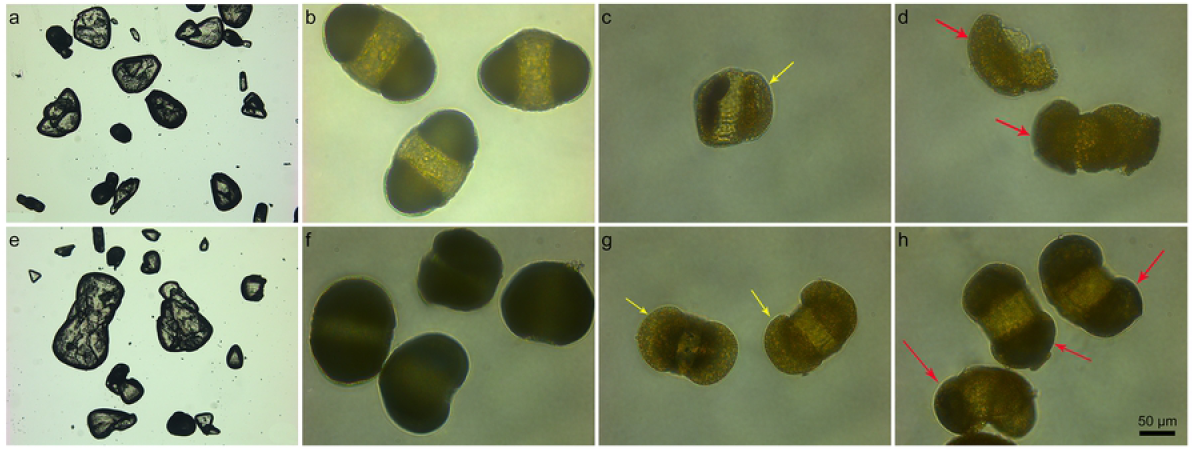
Light microscopy images of a, e) Oxaliplatin drug (*OX), b) Pinus nigra* raw pollen (PN), c) *Pinus nigra* sporopollenin (*PN-SP*), d) Oxaliplatin loaded *Pinus nigra* sporopollenin (*PN-SP-OX*), f) *Cedrus libani* raw pollen (*CD*), g) *Cedrus libani* sporopollenin (*CD-SP*) and h) Oxaliplatin loaded *Cedrus libani* Sporopollenin (*CD-SP-OX*).

**Figure 2.**
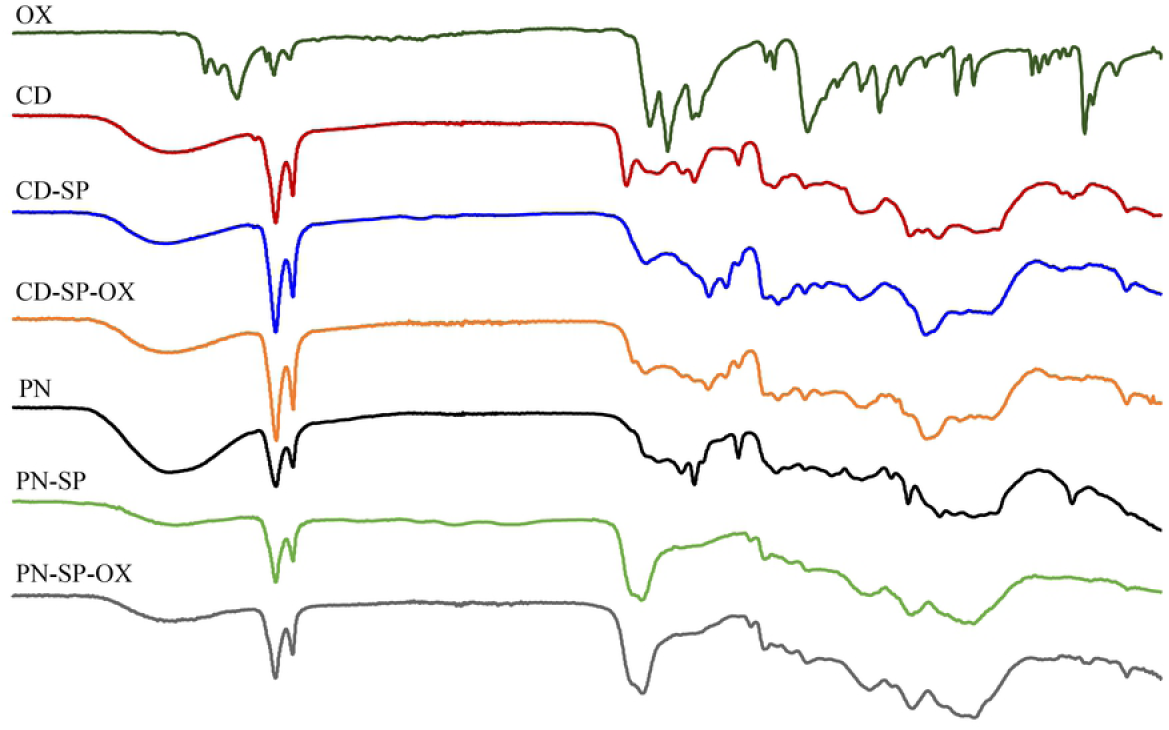
FT-IR spectra of Oxaliplatin drug (*OX), Cedrus libani* raw pollen (*CD*), *Cedrus libani* sporopollenin (*CD-SP*), Oxaliplatin loaded *Cedrus libani* Sporopollenin (*CD-SP-OX*), *Pinus nigra* raw pollen (PN), *Pinus nigra* sporopollenin (*PN-SP*), and Oxaliplatin loaded *Pinus nigra* sporopollenin (*PN-SP-OX*).

**Figure 3.**
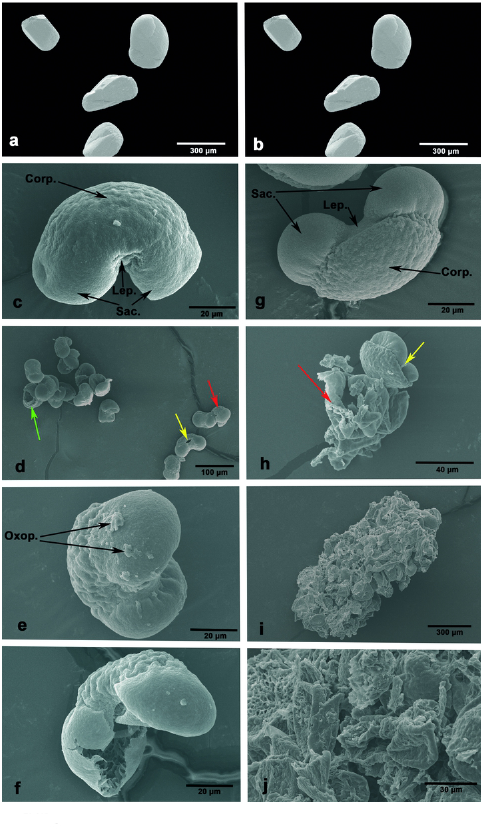
SEM images of a, b) Oxaliplatin drug (*OX);* c) *Cedrus libani* raw pollen (*CD*); d) *Cedrus libani* sporopollenin (*CD-SP*); e, f) Oxaliplatin loaded *Cedrus libani* Sporopollenin (*CD-SP-OX*); *g) Pinus nigra* raw pollen (PN); h) *Pinus nigra* sporopollenin (*PN-SP*) and i, j) Oxaliplatin loaded *Pinus nigra* sporopollenin (*PN-SP-OX*). (The red arrow shows the sporopollenin plaques, the yellow arrows indicate the capsules of the sporopollenin split from the leptoma region, the green arrow shows the capsules of the sporopollenin split from the saccus region).

In the release tests, *PN-SP-OX* initially released 35–40 % of the drug while *CD-SP-OX* revealed a 15–20 % release in the first 4 h. As is clear from SEM images, acid hydrolysis of *CD* retained its structural integrity to a large extent (85–90 %) compared to *PN*. SEM images of *CD-SP-OX* revealed very little superficial Oxaliplatin residues after the encapsulation. This confirmed that the loaded drug to *CD-SP* had been encapsulated inside or adsorbed on the surface of the cages. On the other hand, as was evident from SEM micrographs, the pollens of *P. nigra* lost its structural integrity (90–95%) and formed sporopollenin plaques after treatment with acid. The drug-loaded onto these sporopollenin plaques was adsorbed on the surface and some drug particles remained on the surface and acted as adhesive for other plaques leading to the formation of granule like structures. The drug particles trapped between the sporopollenin plaques and those present superficially were released quickly into the media followed by their slow release from sporopollenin cages and absorption on the walls. These phenomena also explained the initial release in 4 h by *CD-SP-OX* and *PN-SP-OX*.

### Thermogravimetric analysis (TGA)

TGA was carried out to evaluate the thermal stabilities of the pollens and the drug-loaded sporopollenin. The obtained thermograms of *PN* and *CD, PN-SP, CD-SP, OX, PN-SP-OX* and *CD-SP-OX* are given in Fig4. The TG curves reveal that Oxaliplatin has degraded at 270–330°C (Maximum degradation at 283°C). Oxaliplatin was reported to thermally degrade at approximately 300°C in previous studies [34, 35]. During the degradation, elements such as C, H, O, and N were completely degraded leaving only platin residues. The metal residue for Oxaliplatin after TGA degradation was recorded at 46.74% (Fig 4a). The percentage of platin content of Oxaliplatin was calculated at approximately 49.10% and this value was close to the platin residue recorded during thermal degradation. Five different decomposition stages were recorded for the *PN* and *PN-SP* (Fig 4b, d). The initial mass loss in the thermograms of all the tested samples is attributed to the evaporation of adsorbed structural water molecules [36]. The second, third and fourth mass losses can be attributed to the degradation of pollen intine (structural materials such as cellulose, genetic material, lipid, etc.) [37]. Degradation of *PN-SP-OX* was recorded in four stages (Fig 4f). The first decomposition up to 100°C can be attributed to the evaporation of the bound water, while the second decomposition, maximum degradation around 345°C, is due to the degradation of the carbohydrates remaining in the sporopollenin structure after the removal of genetic material and pectin from the pollen (*P. nigra*) structure. The higher ash content in Oxaliplatin-loaded sporopollenin (24.3 %) is due to the platin residue, a structural component of Oxaliplatin. The thermograms of *CD* reveal the thermal degradation in four steps (Fig 4c). The first mass loss around 53.1°C can be ascribed to the evaporation of structural water molecules. The second and third mass losses have occurred at 250 and 380°C, respectively, displaying a big difference. This mass loss is probably caused by the degradation of pectin and cellulose [38]. The maximum thermal decomposition temperatures for pectin and cellulose are 262.8°C and 378.9°C, respectively. The sharp decomposition at 432.5°C may be due to partial depolymerization and disintegration of the sporopollenin wall. For *CD-SP*, three separate degradations were observed (Fig 4e). After the addition of Oxaliplatin (*CD-SP-OX*), the mass loss in the second degradation increased from 37.8% to 41.5% (Fig 4g). The maximum degradation of sporopollenin was achieved at approximately 440°C (Table 2). It is thought that the mass loss occurring around 400–650°C is caused by the deterioration of the wall (exine) of sporopollenin. According to these results, sporopollenin has high thermal stability. Furthermore, it shows that the normal thermal properties of Oxaliplatin change after the incorporation of Oxaliplatin into sporopollenin samples and that Oxaliplatin has high decomposition temperatures. It was noted in previous studies that the higher the thermal decomposition temperature, the greater the thermal stability [39–41]. Mujtaba et al. (2017) reported that *P. orientalis* sporopollenin had greater thermal stability than biopolymers such as cellulose and chitin [17]. In the results of the current study, it was observed that *P. nigra* and *C. libani* sporopollenin had a high dTGmax temperature and this sporopollenin had more stable structures. In this study, the TGA results of *P. nigra* and *C. libani* species did not show a significant difference in terms of thermal stability.

**Figure 4.**
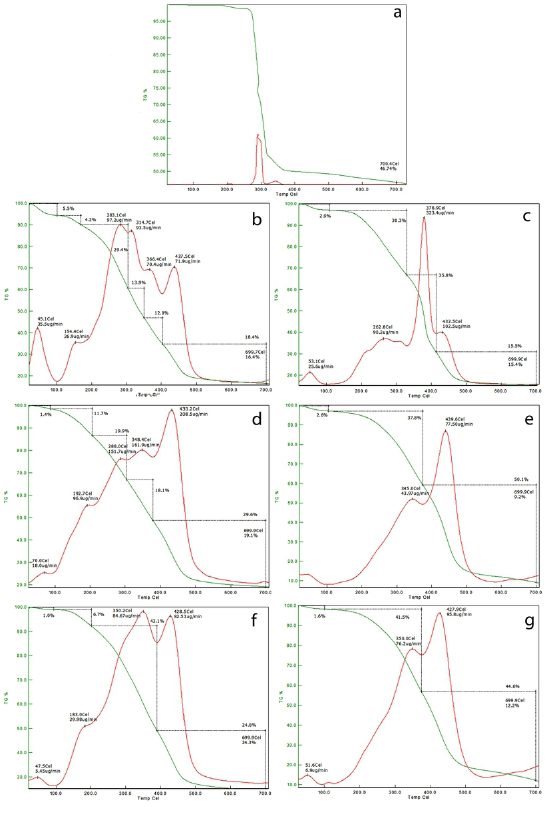
TGA and DTG curves: a) Oxaliplatin drug (*OX), b) Pinus nigra* raw pollen *(PN)*, d) *Pinus nigra* sporopollenin (*PN-SP*), f) Oxaliplatin loaded *Pinus nigra* sporopollenin (*PN-SPOX*), c) *Cedrus libani* raw pollen (*CD*), e) *Cedrus libani* sporopollenin (*CD-SP*) and g) Oxaliplatin loaded *Cedrus libani* Sporopollenin (*CD-SP-OX*).

**Table 2.** Detailed thermogravimetric degradation peaks of Oxaliplatin drug (*OX*), *Pinus nigra* raw pollen (PN), *Pinus nigra* sporopollenin (*PN-SP*), Oxaliplatin loaded *Pinus nigra* sporopollenin (*PN-SP-OX*), *Cedrus libani* raw pollen (*CD*), *Cedrus libani* sporopollenin (*CD-SP*) and Oxaliplatin loaded *Cedrus libani* Sporopollenin (*CD-SP-OX*).

| Samples | Degradation peaks (°C) |  |  |  |  |
| --- | --- | --- | --- | --- | --- |
| *OX* | 283 | - | - | - | - |
| *PN* | 45.1 | 283.1 | 314.7 | 366.4 | 437.5 |
| *PN-SP* | 70.0 | 192.7 | 288.0 | 348.4 | 433.2 |
| *PN-SP-OX* | 47.5 | 183.0 | 350.2 | 428.5 | - |
| *CD* | 53.1 | 262.8 | 378.9 | 432.5 | - |
| *CD-SP* | 55.3 | 345.0 | 439.6 | - | - |
| *CD-SP-OX* | 51.6 | 350.0 | 427.9 | - | - |

### Encapsulation efficiency

The encapsulation efficiency (%) of *CD* and *PN* was recorded as 10.06 % and 38.62% respectively. An important difference was observed among the encapsulation percentages of the two pollens. Here in the current study, a passive drug loading technique was adopted for the Oxaliplatin encapsulation into *CD* and *PN*. The variance in the encapsulation percentages could be attributed to the structural modifications during the isolation. As discussed in detail in the SEM section, acid treatment of *PN* led to major structural disintegration resulting in sporopollenin plaques formation, thus providing a larger surface area for the drug to interact and penetrate. Due to the enhanced surface area, a large amount (38.63%) of Oxaliplatin was absorbed into the sporopollenin porous surface. On the other hand, the *CD* preserved its structural integrity to a large extent during the isolation, thus resulting in sporopollenin cages. As passive loading techniques did not involve any active external forces [42] for loading the drug inside the sporopollenin cages, the loading efficiency of *CD* was recorded lower as compared to *PN*.

**Figure 5.**
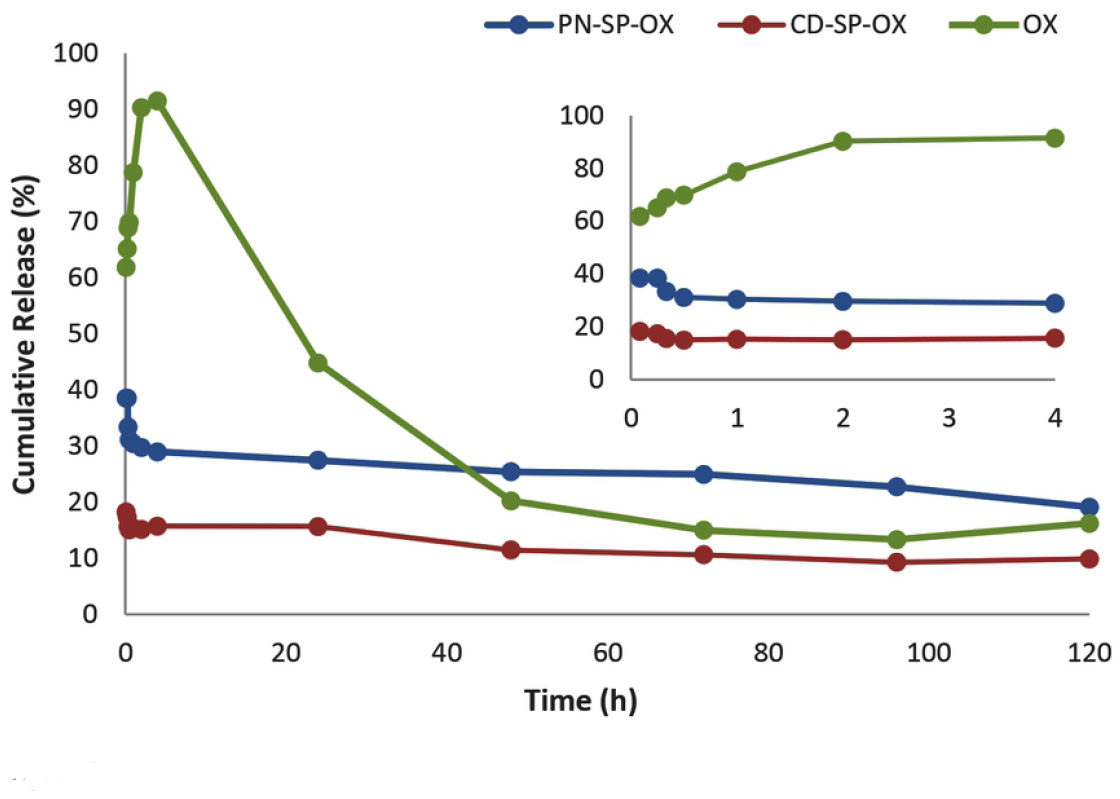
Release properties of sporopollenin microcages from *Cedrus libani* and *Pinus nigra via* passive loading technique in PBS solution.

### Drug release measured in simulated pH solutions

*In vitro* release studies of the Oxaliplatin *(OX)* and Oxaliplatin-loaded *P. nigra* (*PN-SP-OX*) and *C. libani*s poropollenin (*CD-SP-OX*) were performed in PBS (pH 7.4) and results were presented in Fig5.The release profile of the *OX* revealed an initial fast release during the first 4 h and then it dramatically decreased. Overall, drug-loaded sporopollenin showed a slower release rate than the pure drug. The reason behind this slow release could be due to the successful entrapment of the drug inside the sporopollenin microcapsules and plaques of *P. nigra*. Oxaliplatin completely dissolved in PBS in 4 h, whereas in the same time frame the cumulative releases of Oxaliplatin from *CD-SP-OX* and *PN-SP-OX* were recorded at 15.72% and 28.97%, respectively. This fluctuation in the initial release values is attributed to the structural changes (sporopollenin plaques formation) that occur during *P. nigra* sporopollenin extraction process (better explained in SEM section). The results were also found in agreement with the cell studies conducted for checking the release effect in real-time using a real-time cell analyzer (xCELLigence). Also, chitosan nanoparticles [43] and immune hybrid nanoparticles [44] were used as drug delivery systems for Oxaliplatin in the literature. The Oxaliplatin release from chitosan nanoparticles was found to be under 20% for the first 8 h in the Vivek et al. study [43]. Tummala et al. [44] reported that the Oxaliplatin release from immune-hybrid nanoparticles showed a biphasic pattern of drug release at pH=7.4. The authors explained that the cause of the initial burst release was drug molecules that were not entrapped. However, the initial burst release was not observed in the present study. The reason for this result can be due to the entrapment of the drug molecules into the sporopollenin cages and plaques. Consequently, the results of the present in vitro release study demonstrated that *PN-SP* and *CD-SP* could be used as drug carriers.

### Drug release measured *via* xCELLigence RTCA system

In the current study, xCELLigence, a Real-Time Cell Analyzer (RTCA) system, was used to determine the slow-release activity of *P. nigra* and *C. libani*s poropollenin in CaCo-2 and Vero cells in real time (Fig 6). Although traditional drug release and cytotoxic assays are performing well, it is time to support them with advancing technologies of the field. An RTCA system monitors cell growth by measuring electrical impedance, which is represented as a cell index. xCELLigence detects and measures the phenotypic changes occurring in the cells as a result of the release of the drug from sporopollenin through gold microelectrodes mounted at the bottom of the system-specific plates. The prolonged cell proliferation activity reflects the slow release ratio of Oxaliplatin from *PN-SP-OX* and *CD-SP-OX* in Caco-2 and Vero cells and is calculated as normalized cell index by xCELLigence.

**Figure 6.**
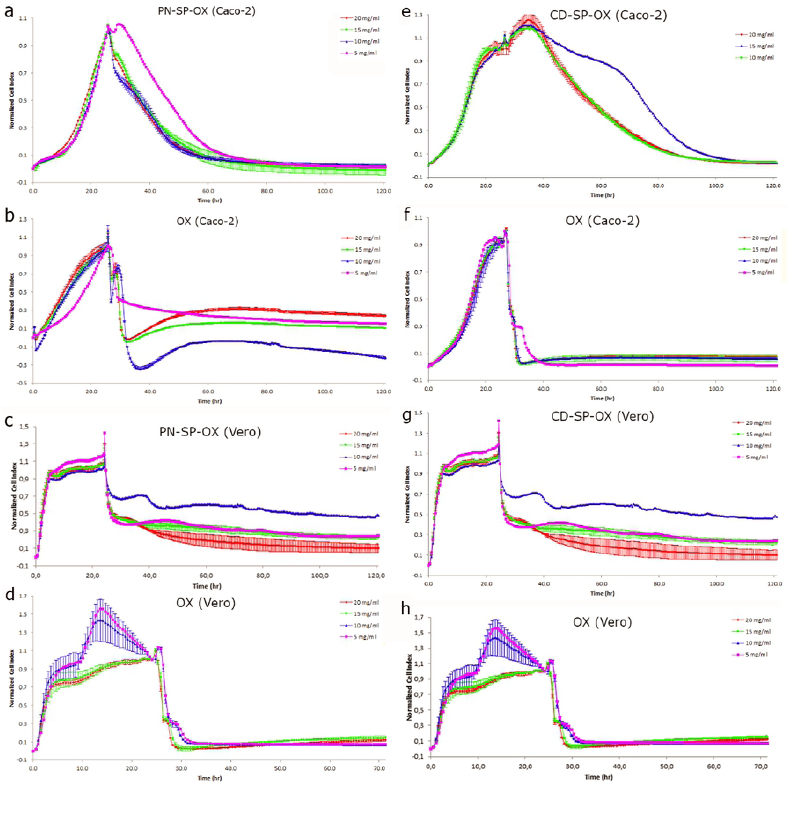
Cell proliferation assay of Caco-2 and Vero cell lines was conducted in the presence of free Oxaliplatin and Sporopollenin loaded with Oxaliplatin using the xCELLigence system. a) sporopollenin of Oxaliplatin loaded *Pinus nigra* sporopollenin/CaCo-2; b) free Oxaliplatin/CaCo-2; c) sporopollenin of Oxaliplatin loaded *Pinus nigra* sporopollenin/Vero, d) free Oxaliplatin/ Vero; e) sporopollenin of Oxaliplatin loaded *Cedrus libani* sporopollenin/CaCo-2; f) free Oxaliplatin/CaCo-2; g) sporopollenin of Oxaliplatin loaded *Cedrus libani* sporopollenin/Vero; h) free Oxaliplatin/ Vero. (pink line: 5 mg/ml; blue line: 10 mg/ml; green line: 15 mg/ml; red line: 20 mg/ml). (The prolonged cell proliferation activity reflects the slow release ratio of Oxaliplatin from *Pinus nigra* and *Cedrus libani* in Caco-2 cells and displayed as normalized cell index by the xCELLigence)

*CD-SP-OX* and *PN-SP-OX* caused cytotoxicity at three doses (5, 10, 15 mg/ml). The IC_50_ values of *PN-SP-OX* were calculated as 10 mg/ml and 14.62 mg/ml for CaCo-2 and Vero cell lines, respectively. The IC_50_ values for *CD-SP-OX* were determined at 5 mg/ml for both examined cell lines. The observed cell index value indicated that the slow release of Oxaliplatin mediated by *CD-SP-OX* and *PN-SP-OX* significantly induced the cell index in a dose-dependent manner on Caco-2 cells compared to Vero cells in over 120 h of incubation (Fig6 a-h). A prolonged-release pattern was observed for *CD-SP-OX* and *PN-SP-OX* compared to free Oxaliplatin drug, with an IC_50_ value of 5 and 14.62 mg/ml after 120 h of incubation. Oxaliplatin displayed an antiproliferative effect in CaCo-2 cells at approximately 5 h at all concentrations, while *PN-SP-OX* revealed a long-delivery effect on CaCo-2 cancer cells for about 41 h. PN-SP-OX showed about an 8 times longer release effect than the control drug (free Oxaliplatin). Interestingly, *PN-SP-OX* applied at a concentration of 10 mg/ml displayed a lower antiproliferative effect on Vero normal cells. This provides a clue that the slow release of the therapeutics also minimizes the harmful effects on normal healthy cells (Fig 6 a, b, c, d). In the case of *CD-SP-OX*, at a concentration of 5 mg/ml, a prolonged proliferation effect on CaCo-2 cells was observed, which lasted for 45 h. At the same concentration, the free Oxaliplatin showed a 5 h antiproliferation activity. Besides, slow-release activity from *CD-SP-OX* was observed in Vero cells (Fig 6 e, f, g, and h).

Considering the overall results, it can be stated that sporopollenin-mediated slow release revealed a prolonged and slow release of Oxaliplatin in both CaCo-2 and Vero cells. But the slow release in healthy cells decreases the antiproliferative effect as a result of the loaded therapeutic i.e., Oxaliplatin. The use of RTCA gives an idea about the beneficial effects of slow release of Oxaliplatin on healthy cells.

### Expression of *FOXO3* and *MYS*

To better understand the apoptosis effects of sporopollenin-mediated slow release of Oxaliplatin on cancer (CaCo-2) and healthy cells (Vero), the expression profiles of two reference genes were investigated for apoptosis by using a qRT-PCR. *FOXO3* and *MYS* genes were selected for this purpose as they were supposed to be responsible for controlling key pathways during the cell apoptosis (detailed functions of both genes have been described in the introduction section).

As shown in Fig 7, a 20-fold increase was recorded for the *FOXO3* mRNA expression in the CaCo-2 cell line when treated with PN-SP-OX (p<0.05) (Fig 7a). Interestingly, in the case of Vero cells, no significant increase was recorded in the expression of FOXO3 after treating it with PN-SP-OX (Fig 7b) (p<0.05).The level of *MYC* gene expression decreased approximately two-fold on the CaCo-2 cell compared to the control (Fig 7c). In a Vero normal cell, the expression level of the *MYC* gene decreased about eight times compared to control (p<0.05) (Fig. 7d). In the case of CD-SP-OX, the level of *FOXO3* gene expression on the CaCo-2 cell increased by about 36 folds compared to the control but a two-fold decrease was detected in Vero cells (p<0.05) (Fig 7e, f). The *MYC* gene expression of CD-SP-OX was not statistically significant compared to the control (OX), but it was an increased gene expression compared to the control of CaCo-2 cells (Fig 7g). The same material showed a three-fold decrease in *MYC* gene expression in Vero cells (p<0.05) (Fig 7h). In summary, the expression of MYC and *FOXO-3* genes increased in CaCo2 cells and decreased in non-cancerous Vero cells. The results demonstrated that the increased expression level of these genes induced apoptosis. The slow release of anticancer drugs in tumor cells may lead to the development of important and novel strategies in the treatment of cancer. However, considering the current clue given by this study, a more detailed investigation is needed into the apoptosis linkage with the slow release on the protein level.

**Figure 7.**
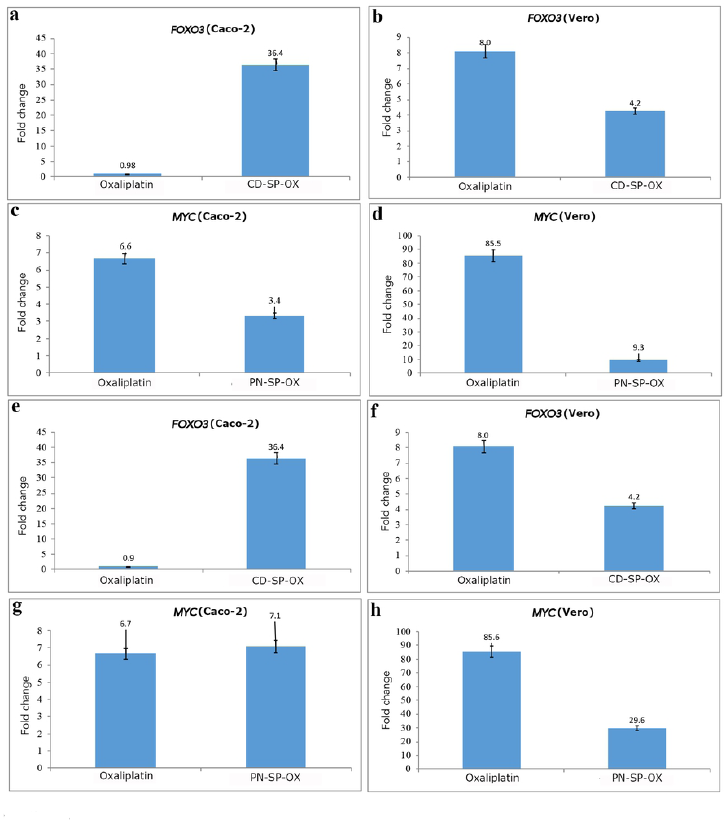
Expression profiles of FOXO3 and MYC genes in Caco-2 and Vero cells treated with Oxaliplatin (OX) and Oxaliplatin-loaded Sporopollenin.

## Conclusions

Sporopollenin exine capsules (SECs) extracted from plant pollens have successfully established its position in the category of biomaterials for control release applications thanks to its robust structural and morphological features, making it an ideal vehicle for encapsulation and control release of drugs. Considering the results of the current study, it can be stated that the sporopollenin of *C. libani* and *P. nigra* successfully demonstrated slow-release abilities of the Oxaliplatin (40–45 h). The sporopollenin-mediated extended release of Oxaliplatin can overcome the negative effect of the routinely used cancer drugs. Both the *in vitro* release assay in PBS and the CaCo-2 as well as Vero cell cultures assay in real-time cell analyzer (xCELLigenece) have confirmed the slow release of *C. libani* and *P. nigra* sporopollenin. Apoptosis plays a critical role in tumorigenesis. In this study, we have provided an understanding of the linkage of sporopollenin-aided slow release and apoptosis mechanisms by checking the expression levels of *MYC* and *FOXO-3* genes. However, further investigation into other genes-related apoptosis pathways is required in order to advance the results of the present study.

## References

1. Agrawal M, Saraf S, Saraf S, Dubey SK, Puri A, Patel RJ, et al. Recent strategies and advances in the fabrication of nano lipid carriers and their application towards brain targeting. Journal of Controlled Release. 2020.

2. Li W, Cao Z, Liu R, Liu L, Li H, Li X, et al. AuNPs as an important inorganic nanoparticle applied in drug carrier systems. Artificial cells, nanomedicine, and biotechnology. 2019;47(1):4222–33.

3. Yang Y, Chen Q, Lin J, Cai Z, Liao G, Wang K, et al. Recent advance in polymer based microspheric systems for controlled protein and peptide delivery. Current medicinal chemistry. 2019;26(13):2285–96.

4. Uthappa U, Kurkuri MD, Kigga M. Nanotechnology Advances for the Development of Various Drug Carriers. Nanobiotechnology in Bioformulations: Springer; 2019. p. 187–224.

5. Duro-Castano A, Talelli M, Rodríguez-Escalona G, Vicent M. Smart Polymeric Nanocarriers for Drug Delivery. Smart Polymers and their Applications: Elsevier; 2019. p. 439–79.

6. Vader P, Mol EA, Pasterkamp G, Schiffelers RM. Extracellular vesicles for drug delivery. Advanced drug delivery reviews. 2016;106:148–56.

7. Soppimath KS, Aminabhavi TM, Kulkarni AR, Rudzinski WE. Biodegradable polymeric nanoparticles as drug delivery devices. Journal of controlled release. 2001;70(1-2):1–20.

8. Stella B, Andreana I, Zonari D, Arpicco S. Pentamidine-loaded Lipid and Polymer Nanocarriers as Tunable Anticancer Drug Delivery Systems. Journal of pharmaceutical sciences. 2019.

9. Madaan K, Kumar S, Poonia N, Lather V, Pandita D. Dendrimers in drug delivery and targeting: Drug-dendrimer interactions and toxicity issues. Journal of pharmacy & bioallied sciences. 2014;6(3):139.

10. Caminade A-M, Turrin C-O. Dendrimers for drug delivery. Journal of Materials Chemistry B. 2014;2(26):4055–66.

11. Tong R, Cheng J. Paclitaxel‐initiated, controlled polymerization of lactide for the formulation of polymeric nanoparticulate delivery vehicles. Angewandte Chemie International Edition. 2008;47(26):4830–4.

12. Yoo HS, Lee KH, Oh JE, Park TG. In vitro and in vivo anti-tumor activities of nanoparticles based on doxorubicin–PLGA conjugates. Journal of controlled Release. 2000;68(3):419–31.

13. Shen Y, Jin E, Zhang B, Murphy CJ, Sui M, Zhao J, et al. Prodrugs forming high drug loading multifunctional nanocapsules for intracellular cancer drug delivery. Journal of the American Chemical Society. 2010;132(12):4259–65.

14. Diego-Taboada A, Maillet L, Banoub JH, Lorch M, Rigby AS, Boa AN, et al. Protein free microcapsules obtained from plant spores as a model for drug delivery: ibuprofen encapsulation, release and taste masking. Journal of Materials Chemistry B. 2013;1(5):707–13.

15. Fan T-F, Hwang Y, Potroz MG, Lau K-L, Tan E-L, Shahrudin Ibrahim M, et al. Degradation of the sporopollenin exine capsules (SECs) in human plasma. Applied Materials Today. 2020;19:100594. doi: https://doi.org/10.1016/j.apmt.2020.100594.

16. Pomelli CS, D’Andrea F, Mezzetta A, Guazzelli L. Exploiting pollen and sporopollenin for the sustainable production of microstructures. New Journal of Chemistry. 2020.

17. Mujtaba M, Sargin I, Akyuz L, Ceter T, Kaya M. Newly isolated sporopollenin microcages from Platanus orientalis pollens as a vehicle for controlled drug delivery. Materials Science and Engineering: C. 2017;77:263–70.

18. Fan T, Park JH, Pham QA, Tan E-L, Mundargi RC, Potroz MG, et al. Extraction of cage-like sporopollenin exine capsules from dandelion pollen grains. Scientific reports. 2018;8(1):1–11.

19. Wang Y, Shang L, Chen G, Shao C, Liu Y, Lu P, et al. Pollen-inspired microparticles with strong adhesion for drug delivery. Applied Materials Today. 2018;13:303–9.

20. Paunov VN, Mackenzie G, Stoyanov SD. Sporopollenin micro-reactors for in-situ preparation, encapsulation and targeted delivery of active components. Journal of Materials Chemistry. 2007;17(7):609–12.

21. Sargin I, Akyuz L, Kaya M, Tan G, Ceter T, Yildirim K, et al. Controlled release and anti-proliferative effect of imatinib mesylate loaded sporopollenin microcapsules extracted from pollens of Betula pendula. International journal of biological macromolecules. 2017;105:749–56.

22. Alshehri SM, Al-Lohedan HA, Al-Farraj E, Alhokbany N, Chaudhary AA, Ahamad T. Macroporous natural capsules extracted from Phoenix dactylifera L. spore and their application in oral drugs delivery. International journal of pharmaceutics. 2016;504(1-2):39–47.

23. Kumar A, Montemagno C, Choi H-J. Smart microparticles with a pH-responsive macropore for targeted oral drug delivery. Scientific reports. 2017;7(1):1–15.

24. Taylor RC, Cullen SP, Martin SJ. Apoptosis: controlled demolition at the cellular level. Nature reviews Molecular cell biology. 2008;9(3):231–41.

25. Brown JM, Attardi LD. The role of apoptosis in cancer development and treatment response. Nature reviews cancer. 2005;5(3):231–7.

26. O’Donnell KA, Wentzel EA, Zeller KI, Dang CV, Mendell JT. c-Myc-regulated microRNAs modulate E2F1 expression. Nature. 2005;435(7043):839–43.

27. Du WW, Fang L, Yang W, Wu N, Awan FM, Yang Z, et al. Induction of tumor apoptosis through a circular RNA enhancing Foxo3 activity. Cell Death & Differentiation. 2017;24(2):357–70.

28. Simpson MG. Plant systematics: Academic press; 2019.

29. Sin A, Pınar N, Mısırlıgil Z, Çeter T, Yıldız A, Alan Ş. Polen Allerjisi (Türkiye Allerjik Bitkilerine Genel Bir Bakış). Ankara: Engin Yayınevi. 2007.

30. Alan Ş, Yıldırım Ö, Pınar N, Seçil D, Keçeli T, Çeter T, et al. Betula pendula Roth (syn= B. verrucosa) polenine duyarlı hastalarda IgE reaktivite profilleri. Asthma Allergy Immunol. 2009;7:100–5.

31. Diego-Taboada A, Beckett S, Atkin S, Mackenzie G. Hollow pollen shells to enhance drug delivery. Pharmaceutics. 2014;6(1):80–96.

32. Domínguez E, Mercado JA, Quesada MA, Heredia A. Pollen sporopollenin: degradation and structural elucidation. Sexual Plant Reproduction. 1999;12(3):171–8.

33. Watson JS, Sephton MA, Sephton SV, Self S, Fraser WT, Lomax BH, et al. Rapid determination of spore chemistry using thermochemolysis gas chromatography-mass spectrometry and micro-Fourier transform infrared spectroscopy. Photochemical & Photobiological Sciences. 2007;6(6):689–94.

34. Zhang D, Zhang J, Jiang K, Li K, Cong Y, Pu S, et al. Preparation, characterisation and antitumour activity of β-, γ-and HP-β-cyclodextrin inclusion complexes of oxaliplatin. Spectrochimica Acta Part A: Molecular and Biomolecular Spectroscopy. 2016;152:501–8.

35. Wu L, Man C, Wang H, Lu X, Ma Q, Cai Y, et al. PEGylated multi-walled carbon nanotubes for encapsulation and sustained release of oxaliplatin. Pharmaceutical research. 2013;30(2):412–23.

36. Tirkistani FA. Thermal analysis of some chitosan Schiff bases. Polymer degradation and stability. 1998;60(1):67–70.

37. Sargın İ Arslan G. Chitosan/sporopollenin microcapsules: Preparation, characterisation and application in heavy metal removal. International journal of biological macromolecules. 2015;75:230–8.

38. Yang H, Yan R, Chen H, Lee DH, Zheng C. Characteristics of hemicellulose, cellulose and lignin pyrolysis. Fuel. 2007;86(12-13):1781–8.

39. Zuluaga-Domínguez C, Serrato-Bermudez J, Quicazán M. Influence of drying-related operations on microbiological, structural and physicochemical aspects for processing of bee-pollen. Engineering in Agriculture, Environment and Food. 2018;11(2):57–64.

40. Mujtaba M, Murat K, Ceter T. An investigation of pollen grain thermal diversity on species level. Communications Faculty of Sciences University of Ankara Series C Biology. 2018;27(2):170–6.

41. Mujtaba M, Murat K, Ceter T. Differentiation of thermal properties of pollens on genus level. Communications Faculty of Sciences University of Ankara Series C Biology. 2018;27(2):177–84.

42. Mundargi RC, Tan E-L, Seo J, Cho N-J. Encapsulation and controlled release formulations of 5-fluorouracil from natural Lycopodium clavatum spores. Journal of Industrial and Engineering Chemistry. 2016;36:102–8.

43. Vivek R, Thangam R, Nipunbabu V, Ponraj T, Kannan S. Oxaliplatin-chitosan nanoparticles induced intrinsic apoptotic signaling pathway: A “smart” drug delivery system to breast cancer cell therapy. International journal of biological macromolecules. 2014;65:289–97.

44. Tummala S, Gowthamarajan K, Satish Kumar M, Wadhwani A. Oxaliplatin immuno hybrid nanoparticles for active targeting: an approach for enhanced apoptotic activity and drug delivery to colorectal tumors. Drug delivery. 2016;23(5):1773–87.

